# Aerobicity stimulon in *Escherichia coli* revealed using multi-scale computational systems biology of adapted respiratory variants

**DOI:** 10.1101/2025.03.13.642450

**Authors:** Arjun Patel, Neha Banwani, Raegan Mink, Divya M. Prabhakaran, Snehal V. Khairnar, Adam M. Feist, Bernhard O. Palsson, Amitesh Anand

## Abstract

Energy homeostasis facilitated by the interplay of substrate-level and oxidative phosphorylation is crucial for bacterial adaptation to diverse substrates and environments. To investigate how bioenergetic systems optimize under restrictive conditions, we evolved ETS variants with distinct proton-pumping efficiencies (1, 2, 3, or 4 proton(s) per electron) on succinate and glycerol. These substrates impose unique metabolic constraints: succinate requires complete gluconeogenesis, while glycerol supports mixed glycolytic and gluconeogenic fluxes. Multi-scale computational analysis of the strains revealed (a) Growth optimization across carbon substrates for multiple ETS variants, (b) A conserved aerobicity stimulon comprising seven independently regulated gene groups that are co-regulated with increasing aerobic capacities, (c) Proteome reallocation linked to aerobicity, validated using genome-scale metabolism and expression modeling, and (d) Carbon source-specific compensatory mutations in succinate transporters and regulatory elements. These findings define the aerobicity stimulon and establish a unifying framework for understanding bacterial respiratory flexibility, demonstrating how transcriptional networks and metabolic systems integrate to achieve energy homeostasis and bioenergetic resilience.

## Introduction

Energy conservation pathways and associated metabolic flexibility are critical for bacterial physiology^1^. The need for the indispensable energy carrier molecule, ATP, is met from several metabolic locations in a cell, with oxidative phosphorylation facilitated by ETS being the primary source. This diversity in energy generation pathways allows bacteria to thrive under various environmental conditions by exploiting substrates that differentially feed into various metabolic reactions. Such flux optimization to achieve robust growth physiology requires an interplay among glycolytic, respiratory, and fermentative pathways.

The pathways’ interplay occurs at a level beyond the classical regulon. A set of gene groups that are co-regulated in response to specific environmental stimuli or stress conditions may not be regulated by a specific regulatory protein. Such co-regulated gene groups are referred to as stimulons. This framework gained traction in the late 20th century, with early studies focusing on bacterial shock responses and stress adaptations^2,3^. However, the enthusiasm surrounding stimulons waned in the early 2000s due to challenges related to clearly defining their boundaries, their overlap with regulons, and the technical limitations of early transcriptomics^4,5^. These difficulties and a shift towards systems biology diminished their relevance.

Recent advancements in RNA-Seq, computational modeling, and integrative omics have rekindled interest in stimulons by providing robust frameworks for their identification and analysis. Modern studies have demonstrated the value of stimulons in understanding adaptive responses to oxidative stress, nutrient deprivation, and other environmental challenges^6–8^. By leveraging these technologies, researchers have begun to uncover novel aspects of bacterial physiology and reestablish stimulons as valuable tools for metabolic research. The cellular bioenergetic network has intricate regulatory features, which are difficult to discern solely based on the biochemical properties or classical regulon concepts. In a genome-wide screening for ATP synthetic activity, genes believed to function synergistically in supporting oxidative phosphorylation were proposed to be regulated differently^9^.

In a recent study, we performed adaptive laboratory evolution of four *Escherichia coli* strains that differed in aerobic ETS composition and, thereby, in the number of proton(s) pumped per electron received from NADH^10^. We observed that regardless of the differences in ETS efficiencies, the variants obtained similar growth rates on a minimal medium with glucose as the carbon source by leveraging flexibility in ATP production reactions. We also observed that the Aero-Type System (ATS), a generalized framework of the ETS with additional connecting bioenergetic pathways that we previously described^10^, maintained a constant proteome composition across *E. coli* strains with differing ETS efficiencies, highlighting its role in robust bioenergetic adaptation. However, such bioenergetic equivalency may be limited to respiro-fermentative metabolites like glucose. Despite glucose being a preferred growth-promoting source, *E. coli* needs to survive on varying carbon sources^11^. *E. coli* can exploit lipids, short-chain fatty acids, amino acids, and nucleotides to meet carbon and energy requirements. Notably, the microbiota produces abundant SCFAs, such as succinate, in the mammalian gut. The chemical nature of the carbon source and the enzymatic composition of the cell determines the flux balance of the catabolic energy generation pathways and anabolic biomass synthetic pathways. In this study, we explore the systems optimization approaches of the ETS variants when supplied with carbon sources that are metabolically more restrictive.

The defect in ETS would be less pronounced on glucose as substrate-level phosphorylation can offset the ATP deficiency. We, therefore, evolved the ETS variants on a non-fermentable carbon source, succinate, to exert strong selection pressure on ETS functioning^12^. We also used a fermentable but energy-poor substrate, glycerol, for a parallel evolution^13^. Glucose undergoes complete glycolysis, while glycerol enters at the midpoint of glycolysis, contributing to both glycolysis and gluconeogenesis. In contrast, succinate is primarily directed towards gluconeogenesis (**Figure 1A-C**). We then performed multi-scale computational analyses on the growth rate-optimized strains. We observed (a) growth optimality on multiple carbon substrates, (b) conservation of proteome allocation to the bioenergetic pathways of the ATS, irrespective of carbon substrate or evolutionary conditions, (c) similar transcriptomic adjustments across seven independent gene groups in all ETS variants studied, revealing an aerobicity stimulon, (d) genome-scale models integrated with omics data identifying the ATS as central to bioenergetic genotype-phenotype relationships, and (e) while glycerol did not require ETS-specific genetic changes, succinate-evolved strains acquired compensatory mutations. This study defines the aerobicity stimulon, providing a novel perspective on how *E. coli* optimally utilizes its ATS across phenotypes. Our findings also demonstrate how modern methodologies can illuminate and redefine classical biological frameworks like stimulons.

**Fig. 1.**
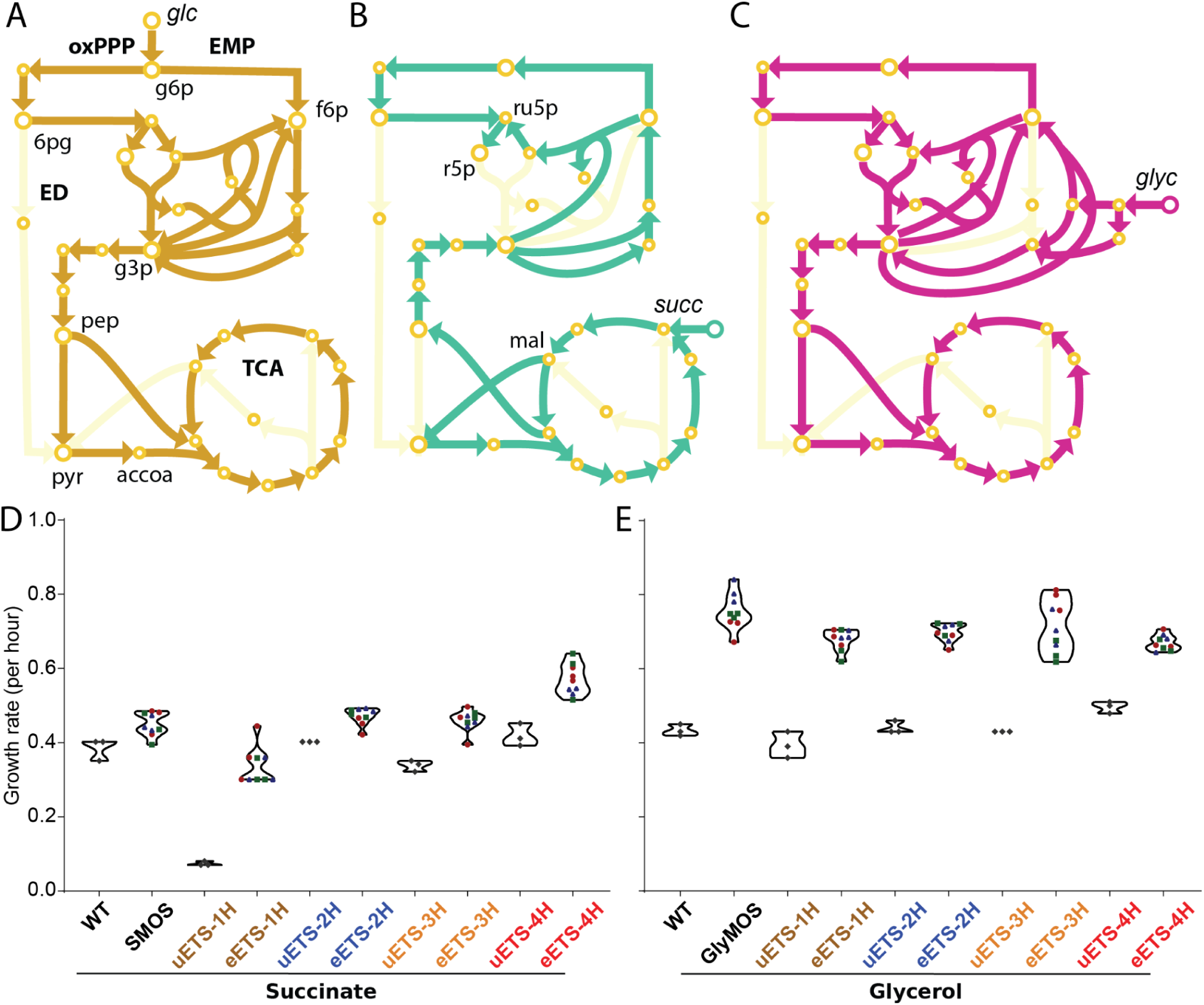
Impact of carbon source on ETS variants. Metabolic flux routing of **(A)** glucose, **(B)** succinate, and **(C)** glycerol in *E. coli* central carbon metabolism. Growth rate of ETS variants on **(D)** succinate and **(E)** glycerol as carbon source before and after growth rate optimization by adaptive laboratory evolution. All laboratory evolutions were performed with three independently evolving lineages. Panels D and E share their y-axis label. The data from the final three flasks of evolution experiments were used to show the growth rates of evolved strains. Different colors represent independently evolved lineages of each strain (Lineage A: red circles, Lineage B: green squares, and Lineage C: blue triangles). [Acronyms-EMP: Embden-Meyerhof-Parnas Pathway, ED: Entner-Doudoroff Pathway, oxPPP: Oxidative Pentose Phosphate Pathway, TCA: Tricarboxylic Acid Cycle, g6p: D-Glucose-6-Phosphate, f6p: D-Fructose-6-Phosphate, 6pg: 6-Phospho-D-Gluconate, g3p: Glyceraldehyde 3-Phosphate, pep: Phosphoenolpyruvate, pyr: Pyruvate, accoa: Acetyl-CoA, ru5p: Ribulose 5-Phosphate, r5p: Ribose 5-Phosphate, mal: Malate, glc: Glucose, succ: Succinate, glyc: Glycerol, WT: Wild Type, SMOS: Succinate Minimal Media Optimized WT Strain, GlyMOS: Glycerol Minimal Media Optimized WT Strain.]

## Results and Discussion

### ETS variants exhibit growth optimization and recover on multiple substrates

The ETS in *E. coli* consists of multiple dehydrogenases and reductases, allowing the bacterium to adapt to aerobic and anaerobic environments^14^. Previous studies have demonstrated a regulatory hierarchy in *E. coli*, where oxygen represses anaerobic pathways, and nitrate further suppresses alternative electron acceptors^14,15^. Despite these controls, the co-expression of multiple respiratory pathways has been shown to enhance flexibility in electron transport^15^. Interestingly, multiple electron flow routes exist for the same electron donor-acceptor pair-NADH to oxygen. Despite the same starting and ending metabolic fate, electrons moving through these alternate routes interact with different redox enzymes. Thus, the alternate routes have distinct proton motive force generation abilities. This allowed us to genetically engineer four distinct ETS variants translocating 1, 2, 3, or 4 protons per electron (designated ETS-nH, where n = 1, 2, 3, or 4)^10^. Building on our prior work with glucose as a carbon source (**Figure 1A**), we extended this framework to investigate the condition-specific performance of the ETS variants under succinate (**Figure 1B**) and glycerol (**Figure 1C**) metabolism—two carbon sources with distinct metabolic and bioenergetic profiles.

Analyzing the growth phenotype of the unevolved strains (called uETS) reveals that the uETS variants grown on succinate exhibited growth rates clearly associated with their H^+^/e^-^ value, with the lowest H^+^/e^-^ strain, uETS-1H, experiencing the most significant growth retardation (**Figure 1D**). This growth pattern confirms the metabolic restriction imposed by non-fermentable carbon sources. In contrast, the glycerol strains showed similar growth rates with no apparent association to their H^+^/e^-^ value (**Figure 1E**). For a detailed systems biology study, we started by performing adaptive laboratory evolution (ALE) of the ETS variants in the two media types in oxic conditions to optimize their growth. The evolved variants, designated as eETS-nHm, were indexed by their replicate endpoints (m = A, B, C). The evolution continued until growth rates stabilized. Similar to the glucose evolutions, despite differences in proton-pumping capacities, all four ETS variants on glycerol ultimately converged on a similar optimized growth rate across replicates (**Figure 1E**), whereas they acquired different optimized growth rates on succinate (**Figure 1D**). ETS-1H, with the lowest ETS efficiency, failed to match the growth rates of the other variants, while ETS-4H acquired the highest growth rate.

Previously, we established a method to categorize *E. coli* phenotypes into aero-types, a quantitative descriptor reflecting cellular respiratory activity and proteome allocation patterns^16^. Aero-types are defined by the multimodal distribution of the proportion of total ATP generated via ATP synthase, influenced by the discrete utilization of ETS enzymes, essentially capturing the aerobicity state of the cell. This classification revealed a non-uniform distribution of growth phenotypes within the rate-yield plane, which could be broadly partitioned into distinct aero-types through sampling simulations. In this study, we applied the aero-type framework to assess the fitness distribution across the ETS variants.

In the previous glucose evolutions, the aerobicity of the evolved strains followed a distinct ranking, with ETS-2H and ETS-4H demonstrating higher aerobic capacities while ETS-3H and ETS-1H exhibiting lower. Specifically, the aerobicity increased as follows: ETS-1H < ETS-3H < ETS-2H < ETS-4H ^10^. Notably, the same hierarchy in aerobicity is preserved when examining the evolved strains grown on succinate (**Figure 2A**) and glycerol (**Figure 2B**), suggesting that the differences in aerobicity are inherently linked to the specific ETS complexes.

**Fig. 2.**
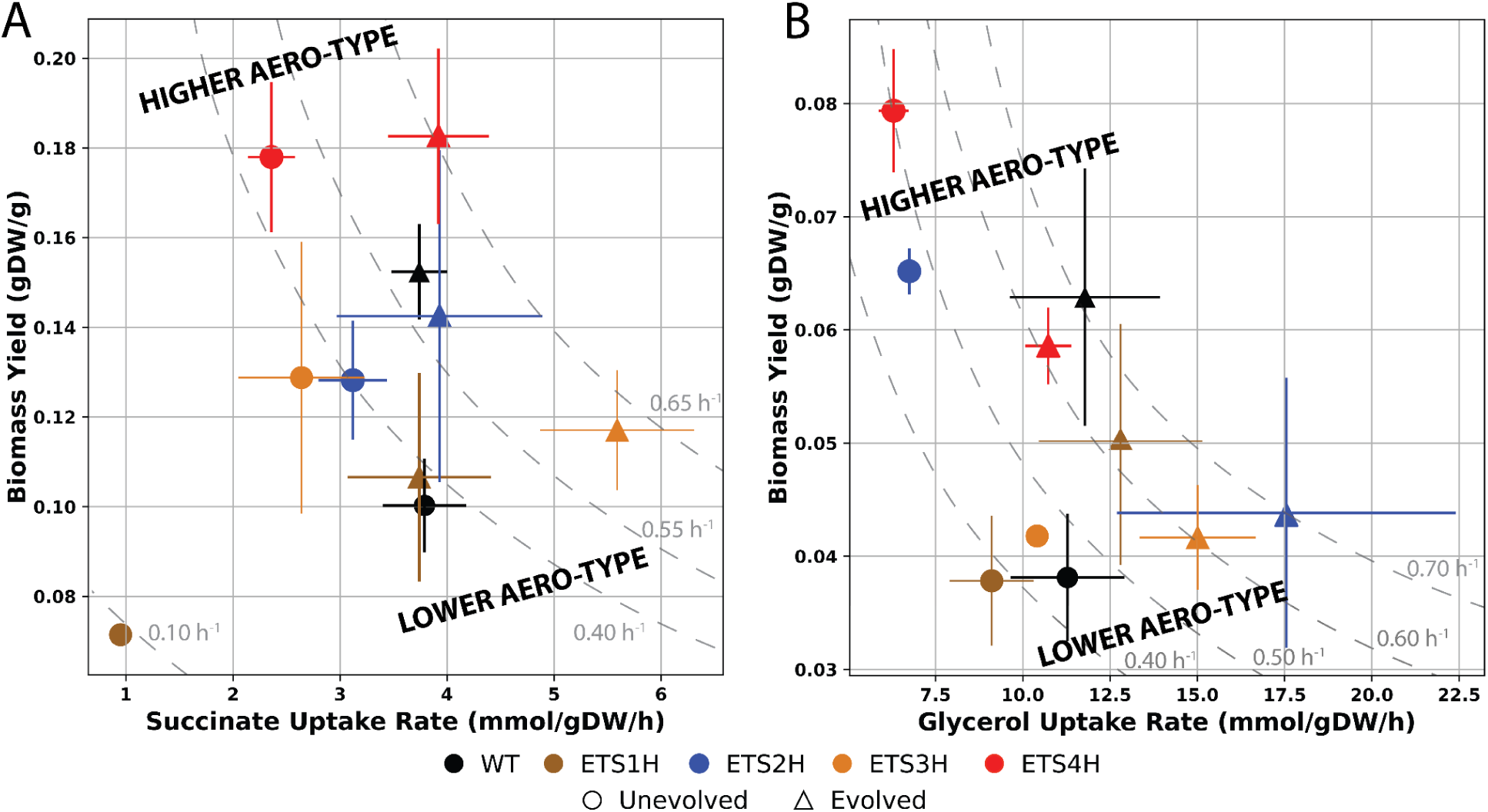
Rate yield planes of ETS variants on each carbon source showcase their different aerobic potentials. Scatter plots of **(A)** succinate and **(B)** glycerol uptake versus biomass yield enable aero-type classification and comparison. Broken gray lines show growth rate isoclines. ETS variants stratify the aero-type spectrum similarly to the strains grown previously on glucose. Source data are provided as the Source Data file.

### Aero-Type System describes convergence in ETS variants’ diverse growth optimization

To understand the observed aerobicity patterns, we employed computational models to validate and further explore the bioenergetic behavior of the ETS variants. Specifically, we used metabolite exchange rates, growth rates, and transcriptomics data as constraints for metabolism and expression models (ME-Models) (Supp. Table 6 and 7, Supp. Data 1) ^17^. These genome-scale metabolic models, which incorporate translation and transcription machinery, allow for the analysis of metabolic flux and proteome allocation^18,19^. By applying this approach, we generated strain-specific ME-Model solutions for each ETS variant grown on succinate and glycerol.

Analysis of quinone fluxes across the ETS variants revealed a clear association between complex usage and aerobicities (**Figure 3A**). The WT strain, which lacks any knockouts, predominantly utilized NDH-I and CYO, a combination that supports high proton-pumping efficiency and is reflective of its high aero-type. In contrast, the ETS variants were constrained by their engineered knockouts, which dictated their complex usage. For instance, ETS-4H and ETS-2H leveraged NDH-I or NDH-II with CYO, reflecting moderately high aero-types. On the other hand, ETS-3H and ETS-1H, which lacked CYO, relied on bd-type oxidases (CYD/CBD) and menaquinones. This aligns with literature indicating that ubiquinone is the primary quinone utilized under aerobic growth conditions, whereas menaquinone predominates under more anaerobic conditions^20^. These observed associations between complex usage and aerobicities highlight how specific combinations of dehydrogenases and oxidases contribute to respiratory efficiency and the bioenergetic state of the cell. These complex usage preferences and their association with aerobicities persist across our previous glucose study, reinforcing the robustness of the observed relationships.

**Fig. 3.**
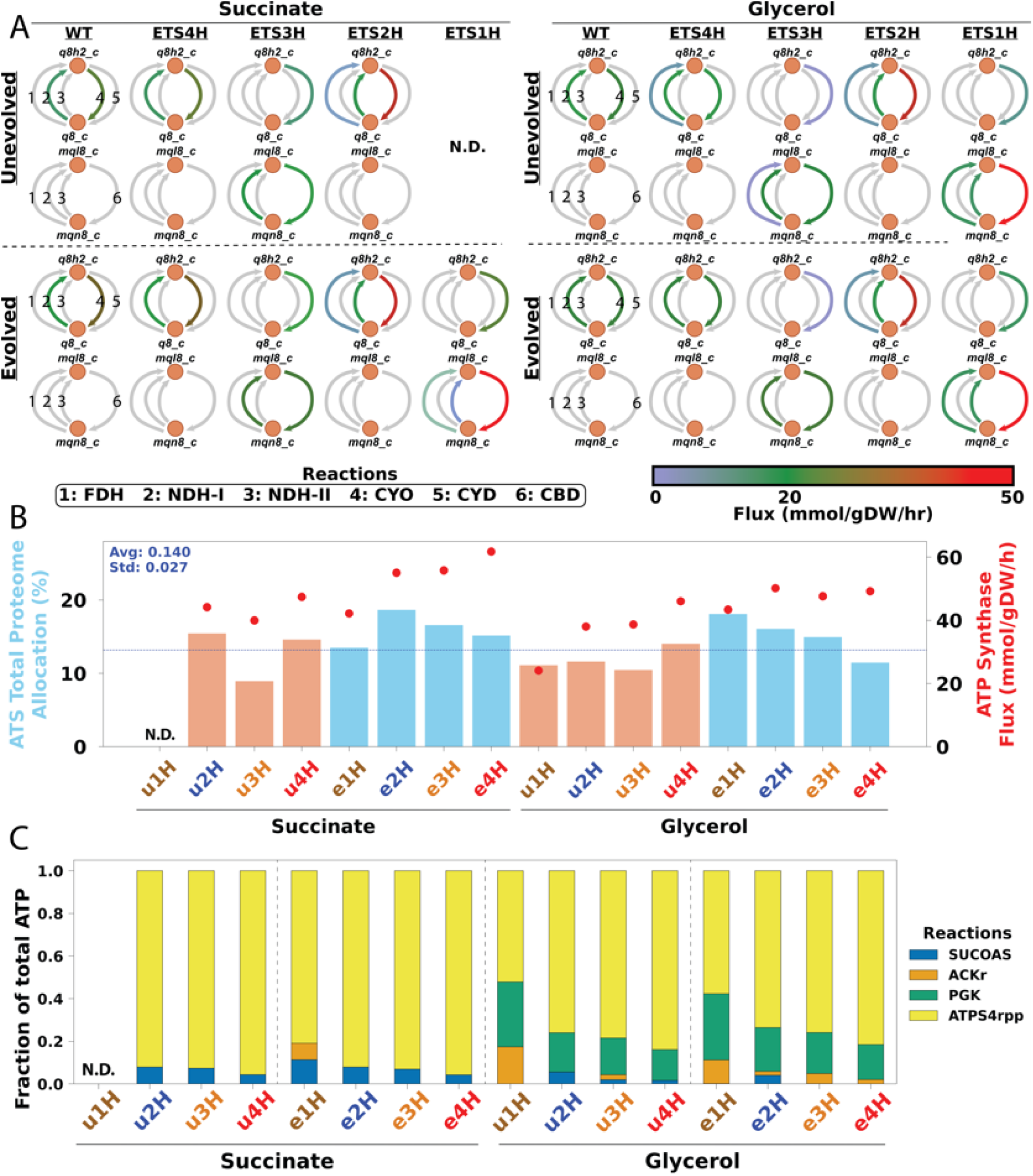
Omics-constrained ME models of ETS variants reveal key characteristics of how strains readjust their bioenergetics. **A)** Quinone flux maps for unevolved and evolved ETS variants. Ubiquinones are in the top loop (q8h2_c and q8_c), while menaquinones are in the bottom loop (mql8_c and mqn8_c). Reactions are labeled 1 through 6 and are Formate Dehydrogenase (FDH), NADH Dehydrogenase I (NDH-I), NADH Dehydrogenase II (NDH-II), Cytochrome bo3 (CYO), Cytochrome bd I (CYD), Cytochrome bd II (CBD), respectively. **B)** Calculated proteome allocation (left y-axis) and flux through ATP Synthase (right y-axis) for all of the ETS variants. **C)** Stacked bar charts of the fraction of total ATP from ATP-producing reactions for all ETS variants. uXH: unevolved X protons per electron ETS variant, eXH: evolved X protons per electron ETS variant, SUCOAS: Succinyl-CoA synthetase, ACKr: Acetate kinase, PGK: Phosphoglycerate kinase, ATPS4rpp: ATP Synthase. Source data are provided as the Source Data file.

Interestingly, NDH-I, despite its molecular mass being approximately 10 times greater than NDH-II^21^, plays a critical role in optimizing ETS efficiency. While NDH-II enables high-turnover electron transfer and alleviates PMF buildup for rapid growth^22^, our findings suggest that NDH-I is more integral for a fully optimized ETS. Similarly, CYO emerged as the most important for ETS efficiency among oxygen reductases, outperforming bd-type reductases CYD and CBD. These preferences for NDH-I and CYO at higher aero-types align with results from a study that modeled 2,200 different phenotype-genotype combinations^16^.

Further analysis revealed the activity of formate dehydrogenases (FDHs) varied with aerobicity. In the unevolved glycerol strains, FDH-O was active in the more aerobic ETS-4H and ETS-2H strains, while FDH-N was utilized in the less aerobic ETS-3H and ETS-1H strains. This is consistent with prior studies showing that FDH-O expression is upregulated under aerobic conditions, whereas FDH-N is induced in anaerobic environments^23,24^.

Analyzing the proteome allocation to the ATS across all strains reveals a consistent distribution, with the models allocating an average of 0.14 proteome fraction (std = 0.027), regardless of growth rate or ATP production via ATP synthase (**Figure 3B**). This finding aligns closely with data from our in-house proteomic expression profiling compendium (n = 162 samples, mean = 0.19, std = 0.027) (**Supplementary Figure 1**)^25^. A similar trend of a consistent proteome was observed in our previous glucose study, further emphasizing the remarkable plasticity of the ATS. Despite this conserved proteome allocation, the strains exhibit different ATP production distributions (**Figure 3C**), with higher aero-type strains relying more heavily on ATP synthase and less on fermentative pathways.

### Transcriptomic adjustments consistency revealing an aerobicity stimulon

We sought to determine whether specific gene groups could explain the shifts in aerobicities, or aero-types, observed across samples. To investigate this, we utilized iModulon analysis and our comprehensive in-house RNA-seq expression profiling compendium (n = 1,035 samples)^26,27^. iModulon analysis employs Independent Component Analysis (ICA), a blind source signal separation algorithm, to decompose RNA-seq expression data into two matrices. The first matrix identifies independently modulated groups of genes, referred to as iModulons (n = 201 iModulons), while the second quantifies the activity levels of each iModulon across all samples in the dataset.

A correlation matrix for all 201 iModulon activities across the 1,035 samples in our compendium was constructed (**Supplementary Figure 2**). Hierarchical clustering of this matrix, followed by thresholding, identified a small number of iModulons with coordinated activities or significant correlations across all of our data. When we overlaid metadata such as the glucose, succinate, and glycerol ETS specialists, along with anaerobic samples from our compendium, we observed that one iModulon cluster distinctly aligned with shifting aerobicities (**Figure 4A**). All scatter plots can be found in **Supplementary Figure 3**. This cluster, which includes seven seemingly unrelated iModulons, emerged as the largest and most robust cluster in the thresholded correlation matrix.

**Fig. 4.**
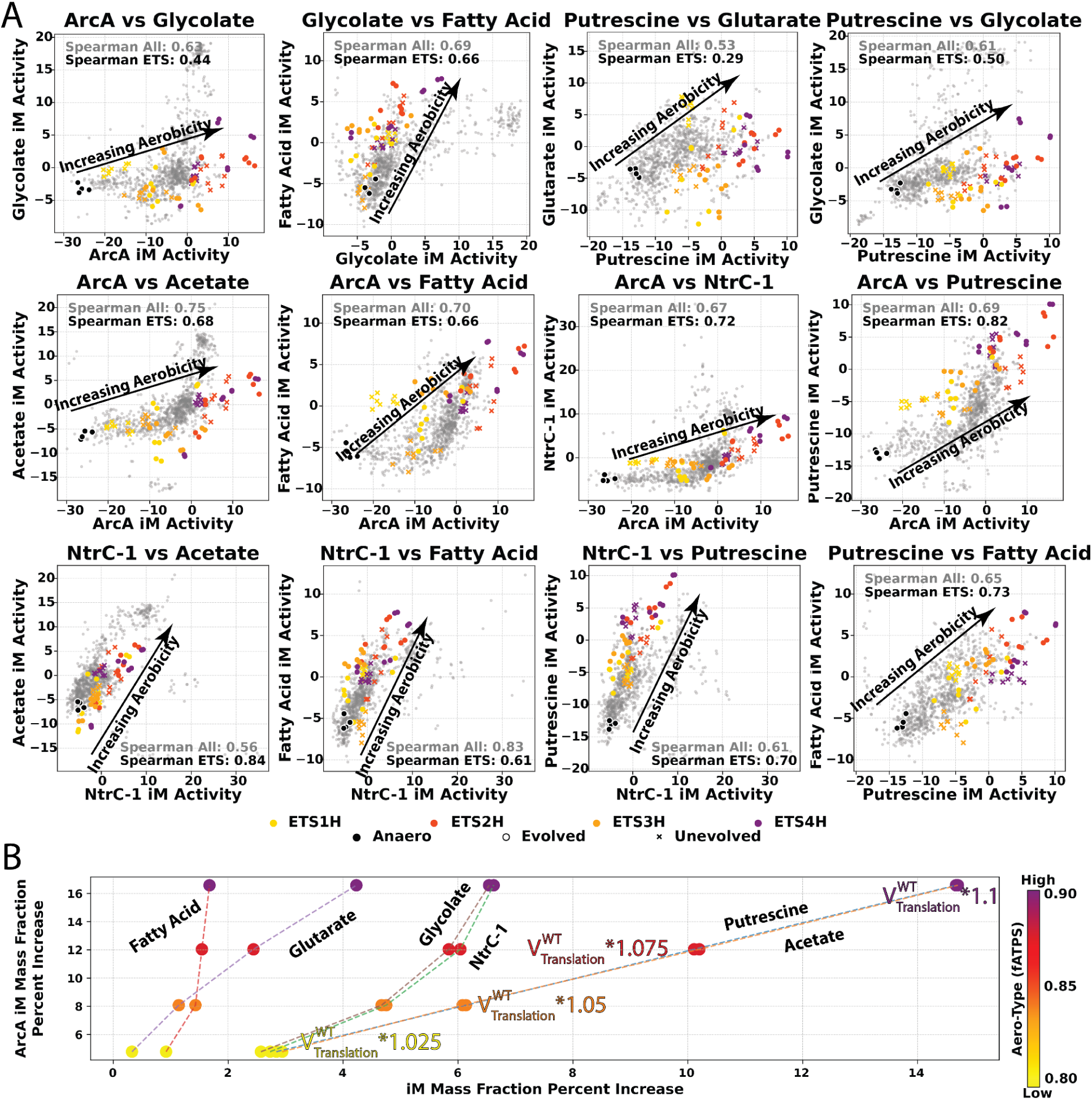
Interoperating multi-omics data and computational biology tools for discovery and validation. **A)** Scatter plots of two iModulon’s Activities from our in-house transcriptome compendium^26,34^ with all glucose, succinate, and glycerol ETS variants labeled. Anaerobic samples are labeled as well to highlight the aerobicity directionality. All other samples in the compendium are colored gray. **B)** Scatter plots of proteome allocation iModulon mass fractions derived from ME Model simulations where the translation flux of ArcA iModulon genes is iteratively increased, starting with the Wild Type flux. Points are colored by the fraction of total ATP generated through ATP synthase (fATPS), which is a calculation of Aero-Type. Source data are provided as the Source Data file.

These seven iModulons‒ArcA, NtrC-1, Acetate, Fatty Acid, Glutarate, Glycolate, and Putrescine‒represent a unified transcriptional response to increasing aerobicities, collectively forming what we term the ***aerobicity stimulon***. Remarkably, the stimulon was consistently observed across all three substrates studied as well as our entire transcriptomic compendium (n = 1035), highlighting its substrate-independent and genotype/phenotype-agnostic role in coordinating the metabolic and bioenergetic adaptations required for scaling aerobicity. The ArcA iModulon, associated with the transcription factor ArcA, encompasses genes involved in aerobic growth, including those encoding ETS complexes^28^. The NtrC-1 iModulon, regulated by NtrC, contains genes from the major arginine degradation pathway in *E. coli*, the arginine N-succinyltransferase (AST) pathway^29^. The Acetate iModulon includes genes for acetate uptake and catabolism, while the Fatty Acid iModulon features the fatty acid degradation (*fad*) genes and is regulated by FadR^30^. The Glutarate iModulon is associated with the 4-aminobutanoate (GABA) transport/catabolism genes *gabDTP*^31^. The Glycolate iModulon comprises the *glcDEFGBA* operon, responsible for glycolate transport and catabolism, and is regulated by the transcription factor GlcC^32^. Lastly, the Putrescine iModulon includes genes for putrescine transport and catabolism, organized in the *puuDRCBE* and *puuAP* operons under the regulator PuuR^33^.

A ME-Model was used to computationally validate the relationships among the seven iModulons. By constraining a wild-type *E. coli* model with increasing levels of translation flux to ArcA iModulon genes, we observed a corresponding increase in proteome allocation to the other six iModulon gene sets (**Figure 4B**). Notably, the Putrescine and Acetate iModulons exhibited coordinated behavior, as did Glycolate and NtrC-1. Furthermore, as these iModulons increased in mass fraction, their aero-type also rose, effectively recapitulating the transcriptomic relationships linked to increasing aerobicities observed in **Figure 4A**.

The underlying biological significance of the aerobicity stimulon is further revealed through its integration with metabolic networks. A network map of these iModulons reveals extensive interconnections and strong links to energy metabolism, revealing their role in effectively scaling aerobicities (**Figure 5**).

**Fig. 5.**
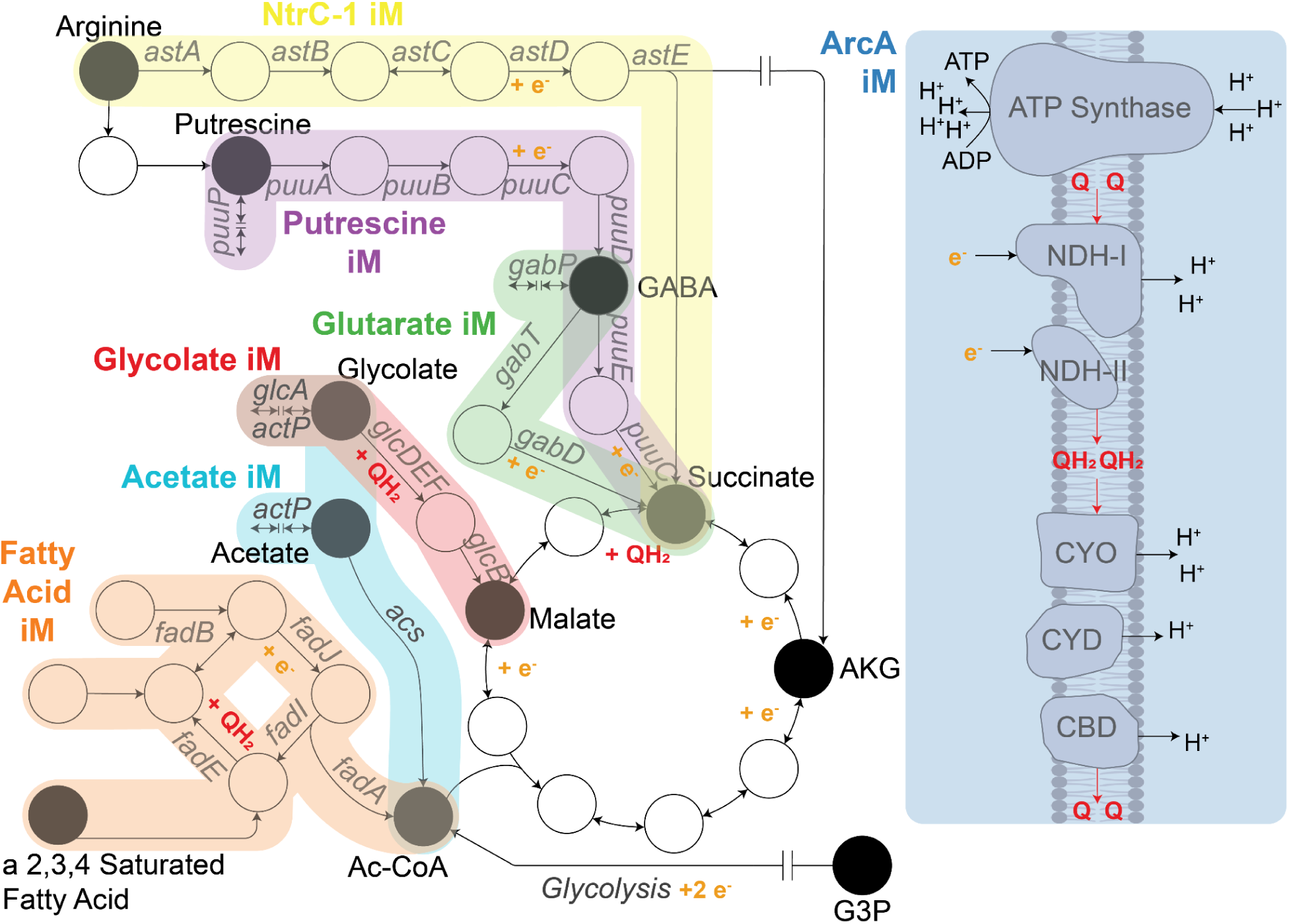
Knowledge-enriched transcriptomic data analytics and computational modeling of metabolic fluxes and proteome allocation elucidate an aerobicity stimulon. A network map of the seven iModulons highlighted in Figure 4 shows their intricate interconnections. These iModulons exhibit a highly coordinated increase in transcriptomic activity as the cell transitions to aerobic conditions. Genes are labeled in gray next to their corresponding arrow. Labeled metabolites are highlighted in black. Quinones are colored in red (QH2: reduced, Q: oxidized), and electron carriers are labeled in orange. NDH-I: NADH Dehydrogenase I, NDH-II: NADH Dehydrogenase II, CYO: Cytochrome bo3, CYD: Cytochrome bd I, CBD: Cytochrome bd II, AKG: 2-oxoglutarate, GABA: 4-aminobutanoate, G3P: Glyceraldehyde 3-Phosphate.

**Fig. 6.**
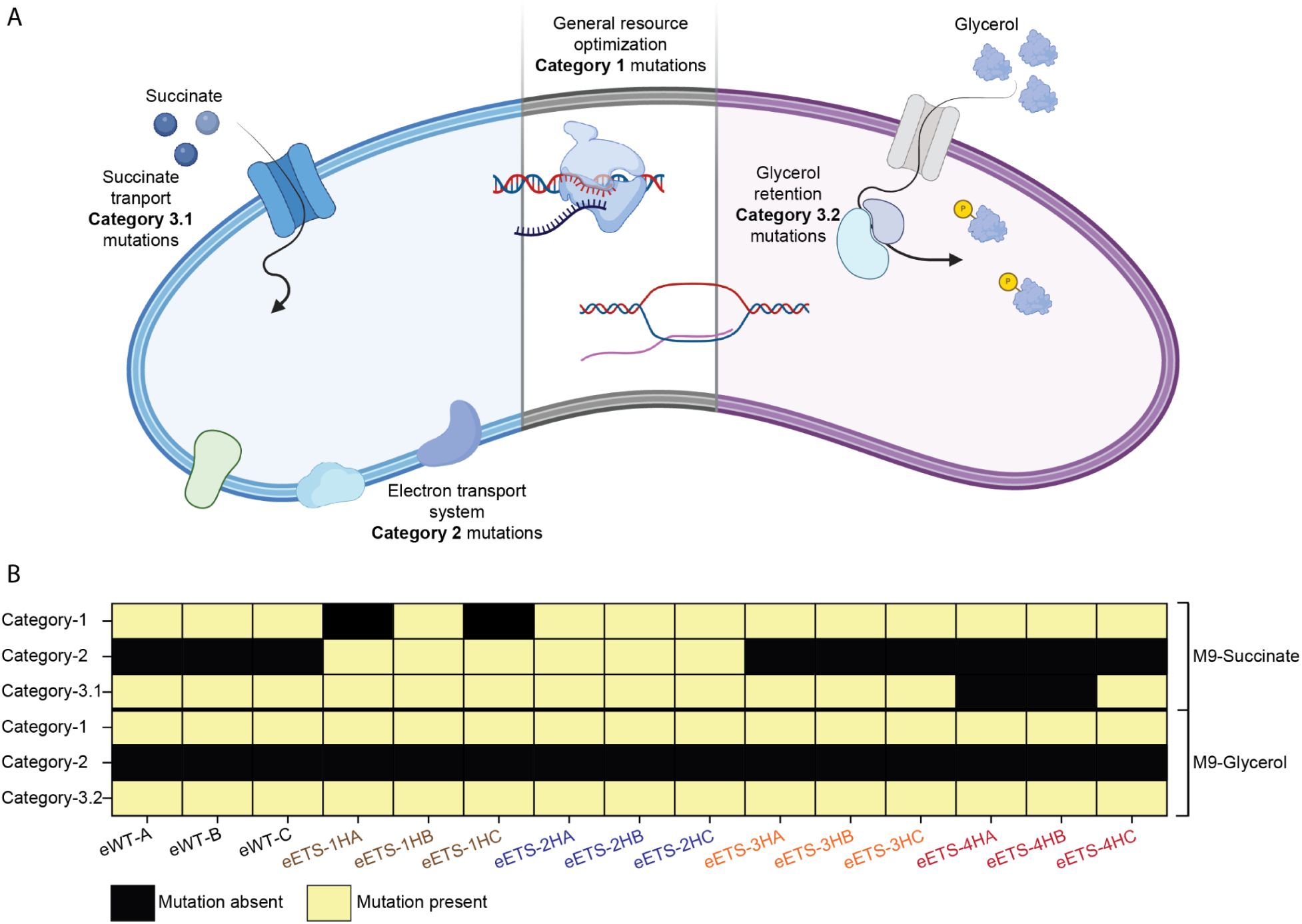
Mutations acquired during growth optimization of the ETS variants on succinate and glycerol. (A) A general scheme for the three major categories of mutations observed. Category 1 mutations are frequently observed during ALE on minimal media and are responsible for supporting growth-supportive cellular resources^35^. Category 2 mutations bring about compensatory transcriptional changes in ETS components. Category 3 mutations influence either carbon source uptake (3.1) or cellular retention (3.2). (B) Heatmap showing the presence or absence of the three major categories of mutations in strains evolved on minimal medium containing either succinate or glycerol as carbon sources. eWT corresponds to SMOS and GlyMOS for the evolution of WT on succinate and glycerol as carbon sources, respectively. Details of the mutations are listed in Supp. Tables 4 and 5.

The metabolic network analysis reveals multiple nodes with high connectivity among the iModulons, including GABA, succinate, and acetyl-CoA. Notably, the acetate/glycolate symporter, *actP*, appears in the Acetate iModulon, while a different symporter, *glcA*, is present in the Glycolate iModulon. Both the Putrescine and Glutarate iModulons degrade GABA using succinate-semialdehyde dehydrogenases (*puuC* and *gabD*, respectively); however, *puuC* generates NADH, whereas *gabD* produces NADPH. The Putrescine and NtrC-1 iModulons connect through agmatine and agmatinase, with Putrescine and Arginine degradation pathways converging to produce succinate. Each iModulon contributes intermediates to the TCA cycle and generates an electron carrier, with the exception of the Acetate iModulon. Furthermore, the Fatty Acid and Glycolate iModulons also produce reduced quinones. Collectively, these intermediates and byproducts enhance the expression of ETS-related genes in the ArcA iModulon, further linking these pathways to aerobic metabolism.

The metabolic complexity and functional interdependencies revealed by this analysis demonstrate that the ETS does not operate in isolation. Instead, it integrates with broader metabolic pathways to enable a systems-level adjustment within the ATS. This alignment of transcriptional responses with respiratory state highlights the critical role of the aerobicity stimulon in bioenergetic optimization.

### Genetic changes enabling growth optimization

Phenotypic gains have an underlying genetic basis. We wanted to examine if specific genetic changes supported the growth improvement of the ETS variants on succinate and glycerol as carbon sources. We observed three categories of mutations in the evolved strains: (1) mutations permitting growth-supportive cellular resource optimization, (2) mutations influencing ETS functioning, and (3) mutations influencing specific carbon-source uptake or retention.

There are several well-characterized mutations known to improve the growth rate of *E. coli* on minimal media. The most commonly observed mutations are related to RNA polymerase and Orotate phosphoribosyltransferase (PyrE). These mutations optimize growth-supportive cellular resources either by global transcriptional changes or by alleviating strain-specific metabolic deficiency ^35,36^. We frequently found fixation of these mutations in the strains evolved on a minimal medium containing either of the carbon sources.

The need to exploit succinate for metabolic needs resulted in the selection of mutations targeting the C4-dicarboxylate transport system^37^. Two common mutations that we observed were in *dctA*, a proton motive force-dependent dicarboxylate transporter, and its transcriptional regulator, *dctR*. Such mutations are reported to improve the succinate transport capacity^38^. We also observed mutations in *dcuB* and *dcuD*, other families of succinate transporters. Interestingly, the WT copy of *dcuD* is believed to be a silent gene due to its transcriptional dormancy^37^. In a previous study, every *E. coli* lineage evolved on a minimal medium containing glycerol acquired mutations in the glycerol kinase gene (*glpK*), the rate-limiting enzyme in glycerol metabolism^39^. We observed similar mutations in *glpK* in every ETS variant evolved on glycerol as the carbon source.

The two ETS variants, ETS-1H and ETS-2H, evolved on succinate acquired mutations influencing the expression of type II NADH dehydrogenase (Ndh; NDH-II)^40^. Notably, these variants have lower ETS efficiency due to the absence of proton-pumping type I NADH dehydrogenase (Nuo; NDH-I). Pyruvate dehydrogenase regulator (PdhR) represses the expression of type II NADH dehydrogenase by binding to the PdhR box upstream of the gene *ndh*. The mutations in this region can alleviate the repression, resulting in the required upregulation of *ndh*. Two eETS-4H strains that evolved on succinate did not acquire category-3.1 mutations, suggesting that these strains’ inherent high ETS efficiency alleviates any stress on cellular energy generation pathways. However, the utilization of glycerol as the carbon source did not impose selection pressure on the ETS.

The findings presented here underscore the remarkable adaptability of the bacterial ETS and its associated metabolism when subjected to environmental and genetic perturbations. Engineered strains with unbranched ETS pathways demonstrated their ability to rewire metabolic networks and achieve near-optimal growth rates by leveraging diverse bioenergetic strategies. The observed plasticity of the ATS across multiple substrates highlights its role as a unifying framework for understanding bacterial respiratory flexibility. By enabling *E. coli* to maintain energy homeostasis through alternate optimal states, the ATS reveals how bacteria adapt their metabolic and respiratory strategies to accommodate specific constraints.

The identification of the aerobicity stimulon adds a novel dimension to this understanding, demonstrating how transcriptional networks contribute to bioenergetic optimization. By combining iModulon analysis with genome-scale metabolic and expression models, this study provides a detailed view of the systems-level adaptations that underpin metabolic resilience. These findings not only deepen our knowledge of bacterial physiology but also establish the ATS as a generalizable descriptor of aerobic bioenergetics, capturing the inherent plasticity and evolutionary versatility of respiration.

## Methods

### Strain generation and adaptive laboratory evolution

The unevolved *Escherichia coli* K12 MG1655 ETS variants used in this study are from the previous project^10,16^. Three independent lineages of each ETS variant were evolved on two separate M9 minimal media preparations, one with glycerol (0.2%) and another with succinate (2 g/l) as the carbon source. Adaptive laboratory evolution was performed as described in previous work^10^.

### DNA sequencing and RNA sequencing

A clone from the evolved strains was used for DNA and RNA sequencing. Total DNA was extracted from an overnight grown culture, and total RNA was extracted from a culture at an A600 ∼0.6. DNA isolation, library preparation, and subsequent analysis were performed as previously described^41^. RNA isolation was performed using the Qiagen RNeasy Mini kit (Cat. No. 74104), following the manufacturer’s protocol of “enzymatic lysis of bacteria.” Ribosomal RNA was removed using the NEBNext rRNA depletion kit Bacteria (Cat. no. NEB E7850X), and cDNA libraries were constructed using the NEBNext Ultra II Directional RNA Library prep kit for Illumina (Cat. no. E7760L).

### Phenotype characterization

Exometabolites were estimated from the culture media. Samples were filtered through a 0.22 µm filter (PVDF, Millipore) and measured using refractive index detection by HPLC (Agilent 1260 Infinity) with a Bio-Rad Aminex HPX87-H ion exclusion column. The HPLC method was the following: injection volume of 10 µL and 5 mM H_2_SO_4_ mobile phase set to a 0.5 mL/min flow rate at 45°C. The phenotype dataset was used for the aero-type classification of the strains as described previously^16^.

### Omics-constrained metabolism and macromolecular expression modeling

The ATS proteome allocation calculation employed here follows the same protocol described in Anand *et al.* 2022^10^. Full details of the procedure, including ATS proteome allocation calculations, ATP generation reaction fluxes, and metabolic flux mapping, can be found in the cited publication. Minor modifications include using succinate or glycerol uptake rate as a constraint rather than glucose uptake rate, and the list of genes constituting the Aero-Type System can be found in Supplementary Data 4. Additionally, the following BiGG metabolic reactions were used for identifying the quinone ETC fluxes^42^: NADH16pp, NADH5, CYTBO3_4pp, CYTBDpp, NADH17pp, NADH10, CYTBD2pp, FDH4pp, and FDH5pp. All flux and proteome mass fraction results can be found in Supplementary Data 3.

### iModulon correlation analysis

iModulon analysis was performed using the iModulon framework ^26,34^. This platform applies Independent Component Analysis (ICA) to decompose RNA-seq data into iModulons, which represent independently regulated gene sets. Activity scores for 201 curated iModulons were calculated across 1,035 transcriptomic samples and are available in our PRECISE-1K compendium^26,34^.

Correlation analysis was conducted on the iModulon activities to identify coordinated independent transcriptional modules. Pairwise Spearman correlations were computed to generate a correlation matrix, which was then hierarchically clustered to identify groups of iModulons with significant co-regulation. Varying thresholding distances of the matrix defined robust clusters, including the seven iModulons that formed the aerobicity stimulon. These clusters were cross-referenced with experimental metadata to assess their relevance to aerobicities. For each iModulon pair within this cluster, activity scores across all samples in the PRECISE-1K compendium were scatter-plotted, with ETS variants grown on the three substrates and anaerobic samples overlaid for comparison. The activity matrix containing all PRECISE-1K samples with the ETS variants on all carbon sources appended at the end can be found in Supplementary Data 2. The anaerobic samples are: p1k_00175, p1k_00176, p1k_00073, and p1k_00074.

### iModulon constrained ME modeling and analysis

ME modeling was used to investigate the impact of increasing fluxes through ArcA iModulon-associated genes on proteome allocation. Simulations were initiated using wild-type (WT) flux values derived from a strain growing with glucose as the carbon source. The fluxes associated with ArcA iModulon genes were progressively increased in increments of 2.5% relative to the WT flux, up to a maximum of 10% of WT flux, in rich media.

For each simulation step (WT, 1.025 * WT, 1.05 * WT, 1.075% * WT, and 1.010 * WT), the ME model calculated the proteome fraction allocated to the seven iModulons comprising the aerobicity stimulon. Total proteome allocation for each iModulon was calculated as follows:

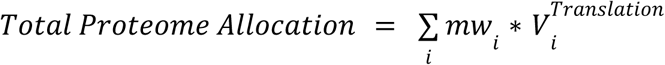

Where *mw*_*i*_ and 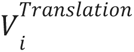 represents the molecular weight and translation flux of the *i*th protein in the iModulon.

## Data Availability

- New transcriptomics accession codes: DNAseq SRA accession number: PRJNA1235627; RNAseq GEO accession number: GSE291717
- Old transcriptomics accession codes: GEO accession number GSE202144 RNAseq TPM data for the new transcriptomics data are available in Supplementary Data 1. The activity matrix for PRECISE-1K and all ETS variants on all carbon sources are available in Supplementary Data 2. Strain-specific ME model calculations for new ETS variants are available in Supplementary Data 3. A list of genes constituting the Aero-Type System is available in Supplementary Data 4. The full PRECISE1k decomposition matrices are available at https://github.com/SBRG/precise1k. Raw RNA-seq data used in PRECISE1k have been deposited at GEO and are publicly available. Accession numbers are listed in the metadata file located in the GitHub repository at the path: data/precise1k/metadata_qc.csv. Source data are provided with this paper.

## Code Availability

All the simulations performed in this manuscript can be reproduced using the FoldME model, which is constructed using the COBRApy toolbox version 0.5.11 for constraint-based modeling and its extension for ME-models, COBRAme version 0.0.9, ECOLIme version 0.0.9, and solveME, all publicly available on Github (https://github.com/SBRG/cobrame, https://github.com/SBRG/ecolime, https://github.com/SBRG/solvemepy)^17,43^. Custom code for constraining and solving ME-models can be found at https://github.com/SBRG/ME-script. Code for our Independent Component Analysis pipeline can be found on GitHub (https://github.com/SBRG/iModulonMiner, https://github.com/SBRG/precise1k), as well as code for all iModulon analysis (https://github.com/SBRG/pymodulon)^44^.

## Acknowledgments

This work was funded by the DAE, India-Tata Institute of Fundamental Research Grant to A.A. and the Novo Nordisk Foundation Grant Numbers NNF10CC1016517 and NNF20CC0035580 to B.O.P. We thank Marc Abrams (Systems Biology Research Group, University of California San Diego) for assistance with paper editing. Icons for some figures were created using our license at Biorender.com

## Author contributions

A.A., and B.O.P. designed the study. N.B., R.M., D.M.P., and A.A. performed experiments. A.P., N.B., S.K., A.M.F., B.O.P., and A.A. analyzed the data. A.P., and A.A. wrote the manuscript with contributions from all other co-authors.

## Competing interests

The authors declare no competing interests.

## Notes

### Competing Interest Statement

The authors have declared no competing interest.

