## Supplementary figures and tables for "Aerobicity stimulon in *Escherichia coli* revealed using multi-scale computational systems biology of adapted respiratory variants"

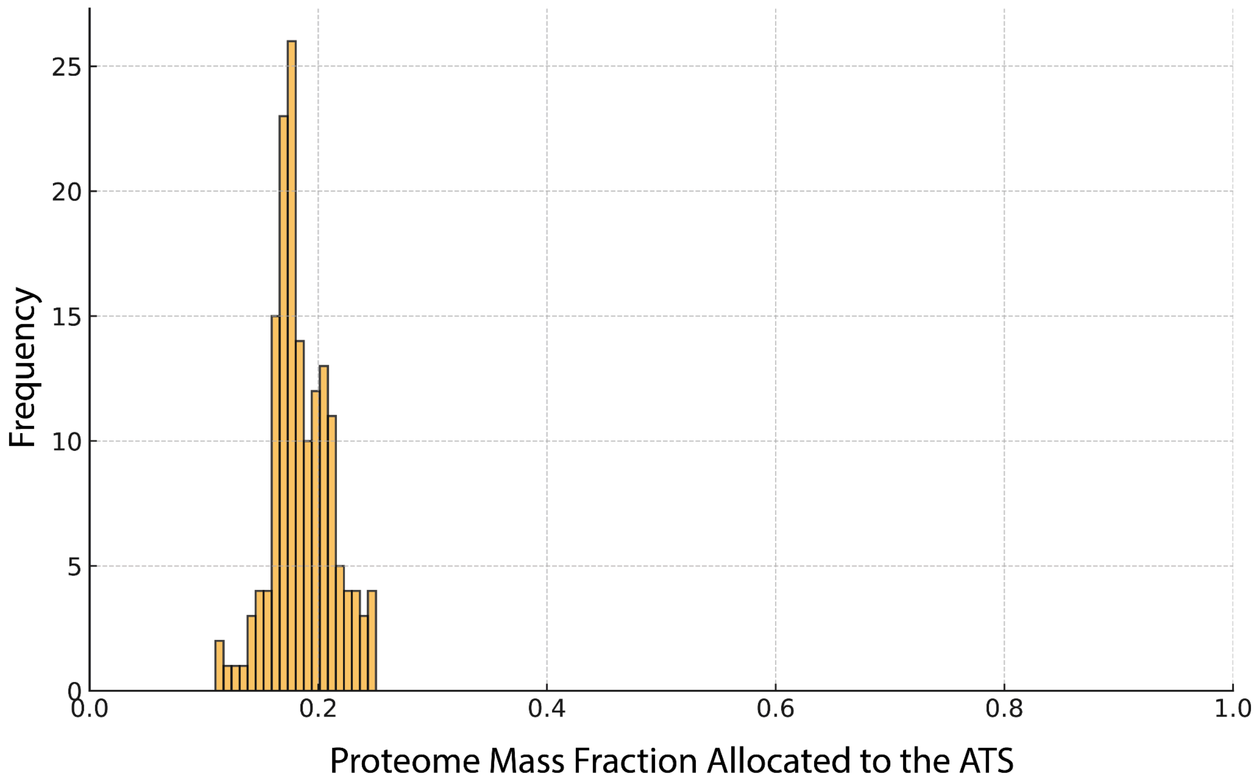


**Supplementary Figure 1. ATS proteome allocation across experimental conditions**

A histogram showing the proteome allocated to the Aero-Type System for all experimental samples from our in-house proteomic expression profiling compendium (n = 162 samples, mean = 0.19, std = 0.027).

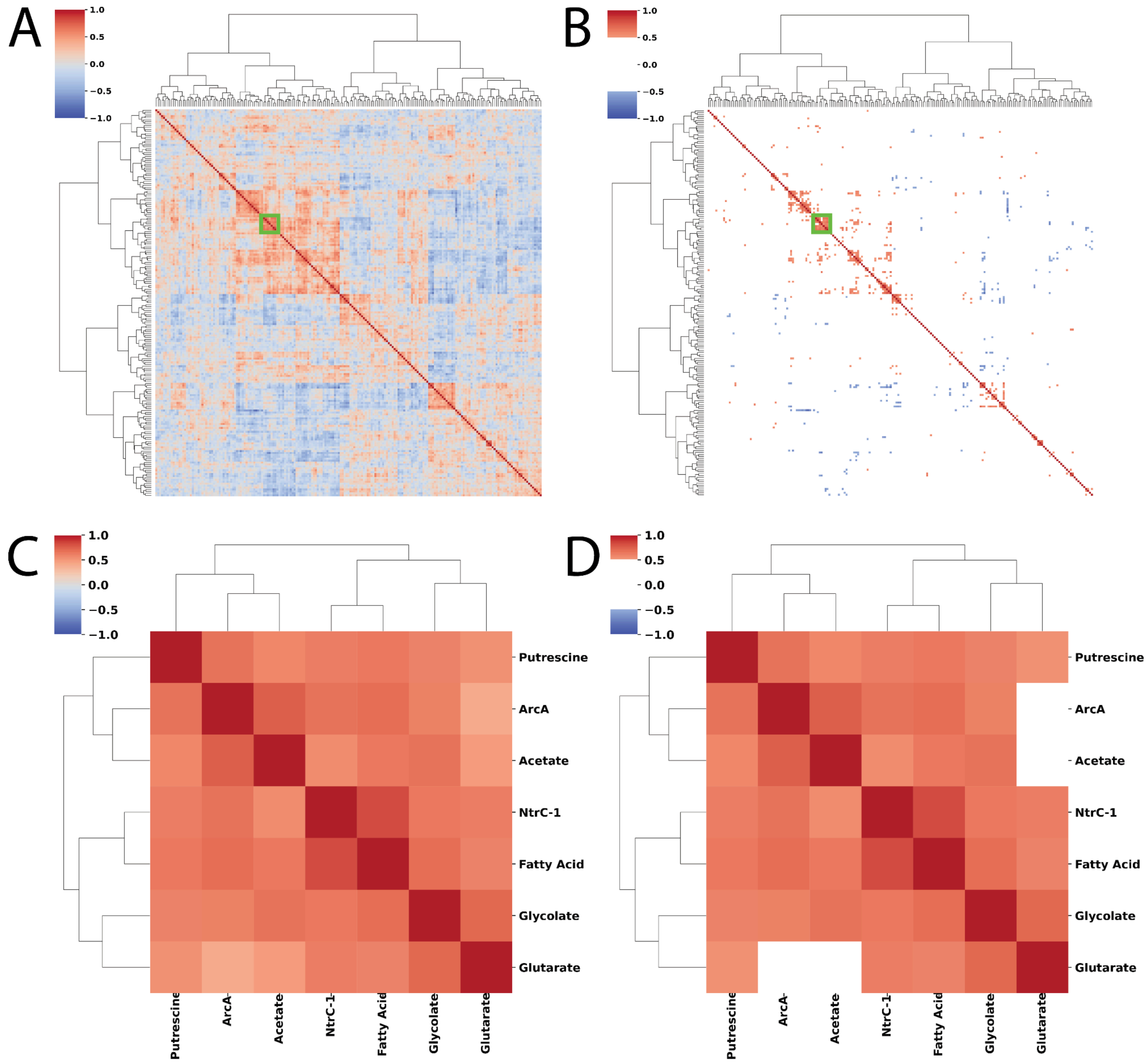


**Supplementary Figure 2. PRECISE-1K activities correlation matrix. A)** Hierarchical clustering heatmap of the correlation matrix for all 201 iModulon activities across 1035 experimental conditions in our in-house transcriptomic compendium, PRECISE-1K. The green box highlights the iModulon cluster that describes aerobic shifts. Applying a correlation threshold of +/- 0.5 **(B)** results in only a few clusters standing out. **C)** A closer look at the seven iModulons in the green box (Putrescine, ArcA, Acetate, NtrC-1, Fatty Acid, Glycolate, Glutarate) that constitute the cluster with and without a threshold **(D)**.

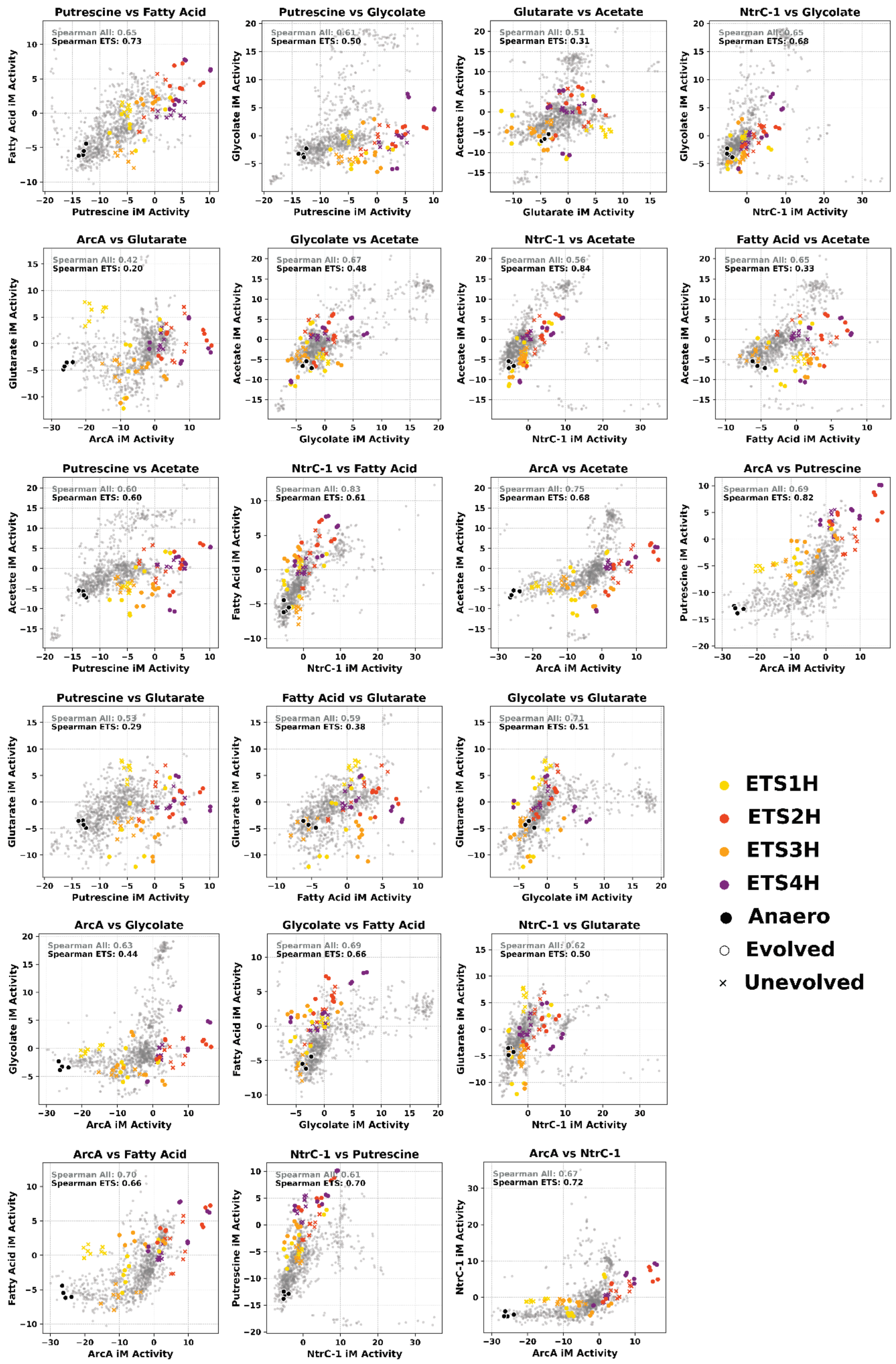
Putrescine vs Fatty Acid
Spearman All: 0.65
Spearman ETS: 0.73
Fatty Acid iM Activity
Putrescine iM Activity
Putrescine vs Glycolate
Spearman All: 0.61
Spearman ETS: 0.50
Glycolate iM Activity
Putrescine iM Activity
Glutarate vs Acetate
Spearman All: 0.51
Spearman ETS: 0.31
Acetate iM Activity
Glutarate iM Activity
NtrC-1 vs Glycolate
Spearman All: 0.65
Spearman ETS: 0.68
Glycolate iM Activity
NtrC-1 iM Activity
ArcA vs Glutarate
Spearman All: 0.42
Spearman ETS: 0.20
Glutarate iM Activity
ArcA iM Activity
Glycolate vs Acetate
Spearman All: 0.67
Spearman ETS: 0.48
Acetate iM Activity
Glycolate iM Activity
NtrC-1 vs Acetate
Spearman All: 0.56
Spearman ETS: 0.84
Acetate iM Activity
NtrC-1 iM Activity
Fatty Acid vs Acetate
Spearman All: 0.65
Spearman ETS: 0.33
Acetate iM Activity
Fatty Acid iM Activity
Putrescine vs Acetate
Spearman All: 0.60
Spearman ETS: 0.60
Acetate iM Activity
Putrescine iM Activity
NtrC-1 vs Fatty Acid
Spearman All: 0.83
Spearman ETS: 0.61
Fatty Acid iM Activity
NtrC-1 iM Activity
ArcA vs Acetate
Spearman All: 0.75
Spearman ETS: 0.68
Acetate iM Activity
ArcA iM Activity
ArcA vs Putrescine
Spearman All: 0.69
Spearman ETS: 0.82
Putrescine iM Activity
ArcA iM Activity
Putrescine vs Glutarate
Spearman All: 0.53
Spearman ETS: 0.29
Glutarate iM Activity
Putrescine iM Activity
Fatty Acid vs Glutarate
Spearman All: 0.59
Spearman ETS: 0.38
Glutarate iM Activity
Fatty Acid iM Activity
Glycolate vs Glutarate
Spearman All: 0.71
Spearman ETS: 0.51
Glutarate iM Activity
Glycolate iM Activity
ArcA vs Glycolate
Spearman All: 0.63
Spearman ETS: 0.44
Glycolate iM Activity
ArcA iM Activity
Glycolate vs Fatty Acid
Spearman All: 0.69
Spearman ETS: 0.66
Fatty Acid iM Activity
Glycolate iM Activity
NtrC-1 vs Glutarate
Spearman All: 0.62
Spearman ETS: 0.50
Glutarate iM Activity
NtrC-1 iM Activity
ArcA vs Fatty Acid
Spearman All: 0.70
Spearman ETS: 0.66
Fatty Acid iM Activity
ArcA iM Activity
NtrC-1 vs Putrescine
Spearman All: 0.61
Spearman ETS: 0.70
Putrescine iM Activity
NtrC-1 iM Activity
ArcA vs NtrC-1
Spearman All: 0.67
Spearman ETS: 0.72
NtrC-1 iM Activity
ArcA iM Activity
ETS1H
ETS2H
ETS3H
ETS4H
Anaero
Evolved
Unevolved

**Supplementary Figure 3. All iModulon pairs in the aerobicity stimulon.**

Scatter plots of iModulon activities for all samples (n=1035) in PRECISE-1K for all pairs of iModulons in the aerobicity stimulon. All glucose, succinate, and glycerol ETS variants are labeled. Anaerobic samples are labeled as well to highlight the aerobicity directionality.

**Supplementary Table 1. Sample code of unevolved strains used in this study**

| Strain | Full ALE code |
|---|---|
| uETS-1H | Δ*nuoB*Δ*cyoB*_A0F0I1R1 |
| uETS-2H | Δ*nuoB*Δ*cydB*Δ*appC*_A0F0I1R1 |
| uETS-3H | Δ*ndh*Δ*cyoB*_A0F0I1R1 |
| uETS-4H | Δ*ndh*Δ*cydB*Δ*appC*_A0F0I1R1 |

**Supplementary Table 2. Sample code of succinate evolved strains used in this study**

| Strain | Full ALE code |
|---|---|
| SMOS-A | A2F102I1R1 |
| SMOS-B | A3F101I1R1 |
| SMOS-C | A4F102I1R1 |
| eETS-1HA | Δ*nuoB*Δ*cyoB*_A7F48I1R1 |
| eETS-1HB | Δ*nuoB*Δ*cyoB*_A5F51I1R1 |
| eETS-1HC | Δ*nuoB*Δ*cyoB*_A6F52I1R1 |
| eETS-2HA | Δ*nuoB*Δ*cydB*Δ*appC*_A9F98I1R1 |
| eETS-2HB | Δ*nuoB*Δ*cydB*Δ*appC*_A11F99I1R1 |
| eETS-2HC | Δ*nuoB*Δ*cydB*Δ*appC*_A10F100I1R1 |
| eETS-3HA | Δ*ndh*Δ*cyoB*_A15F92I1R1 |
| eETS-3HB | Δ*ndh*Δ*cyoB*_A13F94I1R1 |
| eETS-3HC | Δ*ndh*Δ*cyoB*_A16F90I1R1 |
| eETS-4HA | Δ*ndh*Δ*cydB*Δ*appC*_A20F97I1R1 |
| eETS-4HB | Δ*ndh*Δ*cydB*Δ*appC*_A17F92I1R1 |
| eETS-4HC | Δ*ndh*Δ*cydB*Δ*appC*_A18F97I1R1 |

**Supplementary Table 3. Sample code of glycerol evolved strains used in this study**

| Strain | Full ALE code |
|---|---|
| GlyMOS-A | A2F72I1R1 |
| GlyMOS-B | A1F73I1R1 |
| GlyMOS-C | A3F79I1R1 |
| eETS-1HA | Δ*nuoB*Δ*cyoB*_A8F35I1R1 |
| eETS-1HB | Δ*nuoB*Δ*cyoB*_A6F35I1R1 |
| eETS-1HC | Δ*nuoB*Δ*cyoB*_A7F35I1R1 |
| eETS-2HA | Δ*nuoB*Δ*cydB*Δ*appC*_A12F35I1R1 |
| eETS-2HB | Δ*nuoB*Δ*cydB*Δ*appC*_A9F35I1R1 |
| eETS-2HC | Δ*nuoB*Δ*cydB*Δ*appC*_A11F34I1R1 |
| eETS-3HA | Δ*ndh*Δ*cyoB*_A14F32I1R1 |
| eETS-3HB | Δ*ndh*Δ*cyoB*_A13F33I1R1 |
| eETS-3HC | Δ*ndh*Δ*cyoB*_A15F34I1R1 |
| eETS-4HA | Δ*ndh*Δ*cydB*Δ*appC*_A20F65I1R1 |
| eETS-4HB | Δ*ndh*Δ*cydB*Δ*appC*_A18F67I1R1 |
| eETS-4HC | Δ*ndh*Δ*cydB*Δ*appC*_A19F70I1R1 |

**Supplementary Table 4. Key genetic changes in the succinate evolved ETS variants**

| **Strain** | **Category 1** General resource optimization mutations | **Category 2** Electron transport system-related mutations | **Category 3.1** Succinate transport related mutations |
|---|---|---|---|
| SMOS-A | C to T substitution in *rpoC* at genomic position 4188572 | | +15 bp in *dcuB* at genomic position 4348677 |
| SMOS-B | G to T substitution in *rpoC* at genomic position 4189508 | | +5 bp in *dcuB* at genomic position 4348595 |
| SMOS-C | C to T substitution in *rpoC* at genomic position 4189448 | | *dctR*, *dctA* in the duplicated region at genomic position 3560455 |
| eETS-1HA | | C to T substitution in the intergenic region between *ycfP/ndh* at genomic position 1165960 | *dctR*, *dctA* in the duplicated region at genomic position 3653201 |
| eETS-1HB | *rpoD* in the duplicated region at genomic position 3131501 and *pyrE*-*rph* in the amplified region at genomic position 3805701 | C to A substitution in the intergenic region between *ycfP/ndh* at genomic position 1165961 | *dctR*, *dctA* in the duplicated region at genomic position 3584101 and *dcuD* in the amplified region at genomic position 3131501 |
| eETS-1HC | | C to T substitution in the intergenic region between *ycfP/ndh* at genomic position 1165960 | *dctR*, *dctA* in the duplicated region at genomic position 3653201 |
| eETS-2HA | *pyrE*-*rph* in the duplicated region at genomic position 3653201 | G to T substitution in *pdhR* at genomic position 122248 | *dctR*, *dctA* in the duplicated region at genomic position 3619401 |

| | | | |
|---|---|---|---|
| eETS-2HB | G to T substitution in *rpoD* at genomic position 3214754 | C to T substitution in *pdhR* at genomic position 122401 | *dctR*, *dctA* in the duplicated region at genomic position 3619401 |
| eETS-2HC | *rpoA* in the duplicated region at genomic position 3428501 | C to T substitution in the intergenic region between *ycfP/ndh* at genomic position 1165961 | *dctR*, *dctA* in the duplicated region at genomic position 3584201 and *dcuD* in the amplified region at genomic position 3366701 |
| eETS-3HA | Δ1bp in *rpoC* at genomic position 4189434 and *pyrE*-*rph* in the duplicated region at genomic position 3805301 | | *dctR*, *dctA* in the duplicated region at genomic position 3653201 |
| eETS-3HB | A to C substitution in *rpoC* at genomic position 4186605 | | *dctR*, *dctA* in the duplicated region at genomic position 3653201 and C to A substitution in *dcuB* at genomic position 4347763 |
| eETS-3HC | T to G substitution in *rpoB* at genomic position 4183961 | | *dctR*, *dctA* in the duplicated region at genomic position 3652201 |
| eETS-4HA | C to A substitution in *rpoC* at genomic position 4189438 and *pyrE*-*rph* in the duplicated region at genomic position 3721701 | | |

| | | | |
|---|---|---|---|
| eETS-4HB | Δ1 bp in *rpoC* at genomic position 4189451 | | |
| eETS-4HC | G to A substitution in *rpoB* at genomic position 4182880 and *pyrE*-*rph* in the duplicated region at the genomic position 3653201 | | *dctR*, *dctA* in the duplicated region at the genomic position 3653201 |

**Supplementary Table 5. Key genetic changes in the glycerol evolved ETS variants**

| **Strain** | **Category 1** General resource optimization mutations | **Category 2** Electron transport system-related mutations | **Category 3.2** Glycerol retention-related mutations |
|---|---|---|---|
| GlyMOS-A | A to T substitution in *rpoB* at genomic position 4183102 | | T to A substitution in *glpK* at genomic position 4117005 |
| GlyMOS-B | C to A and C to T substitution in *rpoC* at genomic positions 4188572 and 4188929 | | T to G substitution in *glpK* at genomic position 4116250 |
| GlyMOS-C | G to A substitution in *rpoB* at genomic position 4182880 | | G to A substitution in *glpK* at genomic position 4117195 |
| eETS-1HA | G to C substitution in *rpoC* at genomic position 4187603 | | G to C substitution in *glpK* at genomic position 4187603 |
| eETS-1HB | C to T substitution in *rpoB* at genomic position 4182899 | | C to A substitution in *glpK* at genomic position 4117018 |
| eETS-1HC | C to T substitution in *rpoC* at genomic position 4188572 | | C to T substitution in *glpK* at genomic position 4117060 |

| | | | |
|---|---|---|---|
| eETS-2HA | C to A substitution in *rpoC* at genomic position 4187342 | | C to A substitution in *glpK* at genomic position 4116529 |
| eETS-2HB | C to A substitution in *rpoC* at genomic position 4187342 | | C to A substitution in *glpK* at genomic position 4116529 |
| eETS-2HC | A to G substitution in *rpoB* at genomic position 4182881 | | T to A substitution in *glpK* at genomic position 4117005 |
| eETS-3HA | G to T substitution in *rpoB* at genomic position 4183597 | | T to G substitution in *glpK* at genomic position 4116250 |
| eETS-3HB | Δ82 bp in *rph* at genomic position 3815859 | | G to T substitution in *glpK* at genomic position 4116658 |
| eETS-3HC | Δ2 bp in *rph* at genomic position 3815882 | | G to A substitution in *glpK* at genomic position 4117195 |
| eETS-4HA | C to A substitution in *rpoC* at genomic position 4188929 | | Single base substitution in the coding region of *glpK* at genomic position 4116532 |
| eETS-4HB | *hns* and *tdk* in the amplified region at genomic position 1259601 | | C to A substitution in *glpK* at genomic position 4117201 |
| eETS-4HC | A to C substitution in *rpoC* at genomic position 4188864 | | Single base substitution in the coding region of *glpK* at genomic position 4116532 |

**Supplementary Table 6. Phenotypic characterization of succinate evolved ETS variants**

| Strain | Growth rate ($h^{-1}$) | Succinate uptake rate (mmol/gDCW/h) | Acetate secretion rate (mmol/gDCW/h) |
|---|---|---|---|
| WT | 0.35, 0.40, 0.40 | 3.36, 4.14, 3.87 | 3.19, 2.68, 2.15 |
| SMOS | 0.57, 0.62, 0.53 | 3.71, 4.01, 3.49 | ND, 0.55, ND |
| uETS-1H | 0.08, 0.07, 0.07 | 0.92, 0.98, 0.93 | ND, ND, ND |
| eETS-1H | 0.53, 0.51, 0.38 | 3.36, 3.35, 4.52 | 1.40, 1.14, 2.17 |
| uETS-2H | 0.40, 0.40, 0.40 | 2.85, 3.04, 3.47 | 2.97, 3.11, 3.79 |
| eETS-2H | 0.57, 0.58, 0.55 | 2.96, 3.97, 4.87 | ND, ND, ND |
| uETS-3H | 0.35, 0.34, 0.32 | 2.00, 2.76, 3.16 | 1.46, 2.31, 2.16 |
| eETS-3H | 0.53, 0.58, 0.61 | 6.41, 5.02, 5.35 | 2.75, 3.81, 3.73 |
| uETS-4H | 0.45, 0.39, 0.41 | 2.13, 2.56, 2.40 | 0.98, 1.33, 1.33 |
| eETS-4H | 0.60, 0.67, 0.62 | 4.42, 3.47, 3.88 | 3.01, 2.72, 2.60 |

For phenotypic characterization, lineage A has been used. Values for independent biological replicates have been shown in the table. ND: not detected.

Note - uETS variants have been grown in succinate minimal media and used for comparison.

**Supplementary Table 7. Phenotypic characterization of glycerol evolved ETS variants**

| Strain | Growth rate ($h^{-1}$) | Glycerol uptake rate (mmol/gDCW/h) | Acetate secretion rate (mmol/gDCW/h) |
|---|---|---|---|
| WT | 0.45, 0.42, 0.43 | 13.14, 10.63, 10.07 | 0.23, 0.81, 0.26 |
| GlyMOS | 0.67, 0.70, 0.75 | 10.21, 10.90, 14.22 | 3.46, 3.48, 4.08 |
| uETS-1H | 0.39, 0.36, 0.43 | 8.28, 8.56, 10.48 | 1.99, 1.99, 4.88 |
| eETS-1H | 0.82, 0.69, 0.65 | 14.44, 13.85, 10.12 | 7.00, 6.59, 4.79 |
| uETS-2H | 0.43, 0.43, 0.46 | 6.98, 6.57, 6.69 | 0.34, 0.31, ND |
| eETS-2H | 0.72, 0.68, 0.71 | 22.91, 16.28, 13.46 | 3.02, 3.65, 3.20 |
| uETS-3H | 0.43, 0.43, 0.43 | 10.34, 10.55, 10.33 | 2.53, 2.29, 2.13 |
| eETS-3H | 0.63, 0.62, 0.60 | 16.91, 14.31, 13.82 | 7.09, 6.27, 5.90 |
| uETS-4H | 0.51, 0.50, 0.48 | 6.77, 6.19, 5.92 | ND, ND, ND |
| eETS-4H | 0.58, 0.64, 0.55 | 11.41, 10.69, 10.08 | 2.86, 2.62, 2.44 |

For phenotypic characterization, lineage A has been used. Values for independent biological replicates have been shown in the table. ND: not detected.

Note - uETS variants have been grown in glycerol minimal media and used for comparison.
